# Functional Comparison to Ezh2 Reveals PRC2-Independent Functions of Jarid2 in Hematopoietic Stem Cell Lineage Commitment

**DOI:** 10.1101/2025.07.18.665536

**Authors:** Hassan Bjeije, Wentao Han, Nancy Issa, Aishwarya Krishnan, Infencia Xavier Raj, Jason Arand, Yanan Li, Wei Yang, Jeffrey A. Magee, Grant A. Challen

**Affiliations:** Division of Oncology, Department of Medicine, Washington University School of Medicine, St. Louis, MO, USA, 63110; Department of Pediatrics, Washington University School of Medicine, St. Louis, MO, USA, 63110

## Abstract

Previous studies showed the Polycomb Repressive Complex 2 (PRC2) co-factor Jarid2 represses self-renewal transcriptional networks in mouse multipotent progenitor cells (MPPs). But only a fraction of de-repressed HSC-specific genes were associated with loss of H3K27me3, implying Jarid2 may have non-canonical (PRC2-indpendent) in hematopoiesis. Here we sought to delineate any PRC2-independnent functions by comparing stem and progenitor cells genetically deficient for either *Jarid2* or *Ezh2* (enzymatic component of PRC2). Loss of *Ezh2* increased myeloid differentiation in transplantation assays. In contrast, loss of *Jarid2* enhanced T-cell output. Single cell transcriptomics showed while loss of *Jarid2* had minimal impact across progenitor populations, loss of *Ezh2* led to accumulation of lymphoid-biased MPP4 cells and B-cell progenitors in the bone marrow. Functional assays confirmed a differentiation block at the pre-pro B-cell stage. The maturational arrest of *Ezh2*-deficient B-cell progenitors contrasts with increased T-cell output from loss of *Jarid2*, suggesting *Jarid2* has non-canonical functions in hematopoiesis.

## INTRODUCTION

Hematopoietic stem cells (HSCs) maintain blood homeostasis through the balance of self-renewal to sustain the HSC pool and differentiation that gives rise to all mature blood cell lineages (Orkin and Zon, 2008; Wilson and Trumpp, 2006). This delicate balance is governed by networks of transcriptional and epigenetic regulators (Buenrostro et al., 2018; Lara-Astiaso et al., 2014). Among the most critical epigenetic regulators is the Polycomb Repressive Complex 2 (PRC2), a histone methyltransferase complex that catalyzes trimethylation of histone H3 on lysine 27 (H3K27me3), a modification associated with transcriptional repression. The enzymatic subunit of PRC2 is Enhancer of Zeste Homolog 2 (Ezh2) which catalyzes the methylation reaction (Margueron and Reinberg, 2011; Simon and Kingston, 2009). Previous studies have demonstrated that *Ezh2* is essential for maintenance of HSC identity and function. Conditional deletion of *Ezh2* in the hematopoietic system results in loss of HSC quiescence, premature differentiation and stem cell exhaustion (Mochizuki-Kashio et al., 2011; Xie et al., 2014). The importance of PRC2 in regulating hematopoiesis is underscored by the prevalence of *EZH2* mutations in a range of blood cancers (Morin et al., 2010; Nikoloski et al., 2010; Score et al., 2012).

PRC2 does not intrinsically possess genome binding selectivity. Locus specificity is provided by the binding of different accessory co-factors that recruit this complex to distinct genomic sites in different cell types (Kasinath et al., 2021; Pasini et al., 2010; Sarma et al., 2008). PRC2 is divided into two subcomplexes defined by the co-factor binding: PRC2.1 (PHF1, MTF2, PHF19) and PRC2.2 (JARID2, AEBP2)(Glancy et al., 2023; Healy et al., 2019; Loh et al., 2021). These variants modulate PRC2 activity, chromatin targeting, and context-specific gene regulation during development and stem cell fate decisions (Kloet et al., 2016; Peng et al., 2009). Our previous studies highlighted a critical role for the PRC2.2 accessory protein Jarid2 in regulation of hematopoiesis. *Jarid2* is essential for repression of self-renewal transcriptional networks in mouse multipotent progenitor cells (MPPs). Conditional deletion of *Jarid2* results in de-repression of HSC-specific genes in MPPs which conveys ectopic self-renewal potential to this normally transiently-engrafting cell population (Celik et al., 2018). But molecular analysis revealed that only a fraction of the de-repressed HSC-specific genes were associated with loss of H3K27me3 in *Jarid2*-null MPPs (Celik *et al*., 2018). This implies Jarid2 may have some non-canonical (PRC2-independent) activities that regulate HSC fate.

Here we sought to delineate PRC2-independent functions of Jarid2 by comparing the functional and molecular properties of stem and progenitor cells genetically deficient for either *Jarid2* or *Ezh2*. Loss of *Ezh2* leads to myeloid skewing and peripheral lymphoid deficiency in competitive transplantation assays. In contrast, loss of *Jarid2* enhanced peripheral blood engraftment with particular augmentation of T-cell output. Single cell transcriptomics showed while loss of *Jarid2* had minimal impact across early progenitor populations, loss of *Ezh2* led to accumulation of lymphoid-biased MPP4 cells in the bone marrow distinguished by increased Smad7-mediated TGFβ signaling repression and spurious expression of HSC-specific genes. CITE-seq analysis further revealed an increase in early B-cell progenitors in *Ezh2*-deficient bone marrow with functional assays showing a differentiation block at the pre-pro B-cell stage. Cumulatively, our data show that Ezh2 restricts myeloid potential and is necessary for lymphoid priming in HSCs, whereas Jarid2 restricts T-cell differentiation. The contrasting differences between stem and progenitor cells lacking *Ezh2* versus *Jarid2* strongly suggest that Jarid2 has non-canonical functions that regulate HSC fate and lineage specification.

## RESULTS

### Genetic Deletion of Both *Jarid2* and *Ezh2* Leads to Depletion of Phenotypically-Defined HSCs

To compare the functional consequences of *Jarid2* and *Ezh2* loss-of-function, we generated inducible conditional knockout mouse models by crossing *Jarid2*^fl/fl^ and *Ezh2*^fl/fl^ mice to the Mx1-Cre strain. Recombination of floxed alleles (=”Δ”) was induced in eight-week-old mice by injection of polyinosinic:polycytidylic acid (pIpC). Successful gene knockout was confirmed by PCR (**Fig. 1A**) and was highly efficient in hematopoietic stem cells (HSCs; **Fig. 1B**). Immunophenotypic analysis (**Fig. 1C**) revealed no major differences in the abundance of most primitive hematopoietic stem and progenitor cell (HSPC) populations, except that both *Jarid2*^Δ/Δ^ and *Ezh2*^Δ/Δ^ mice showed depletion of phenotypically-defined (Lineage-Sca-1+ c-Kit+ Flk2-CD48-CD150+) HSCs (**Fig. 1D**). These genetic mouse models provide powerful tools to compare the functional roles of Ezh2 and Jarid2 in HSPCs.

**Figure 1:**
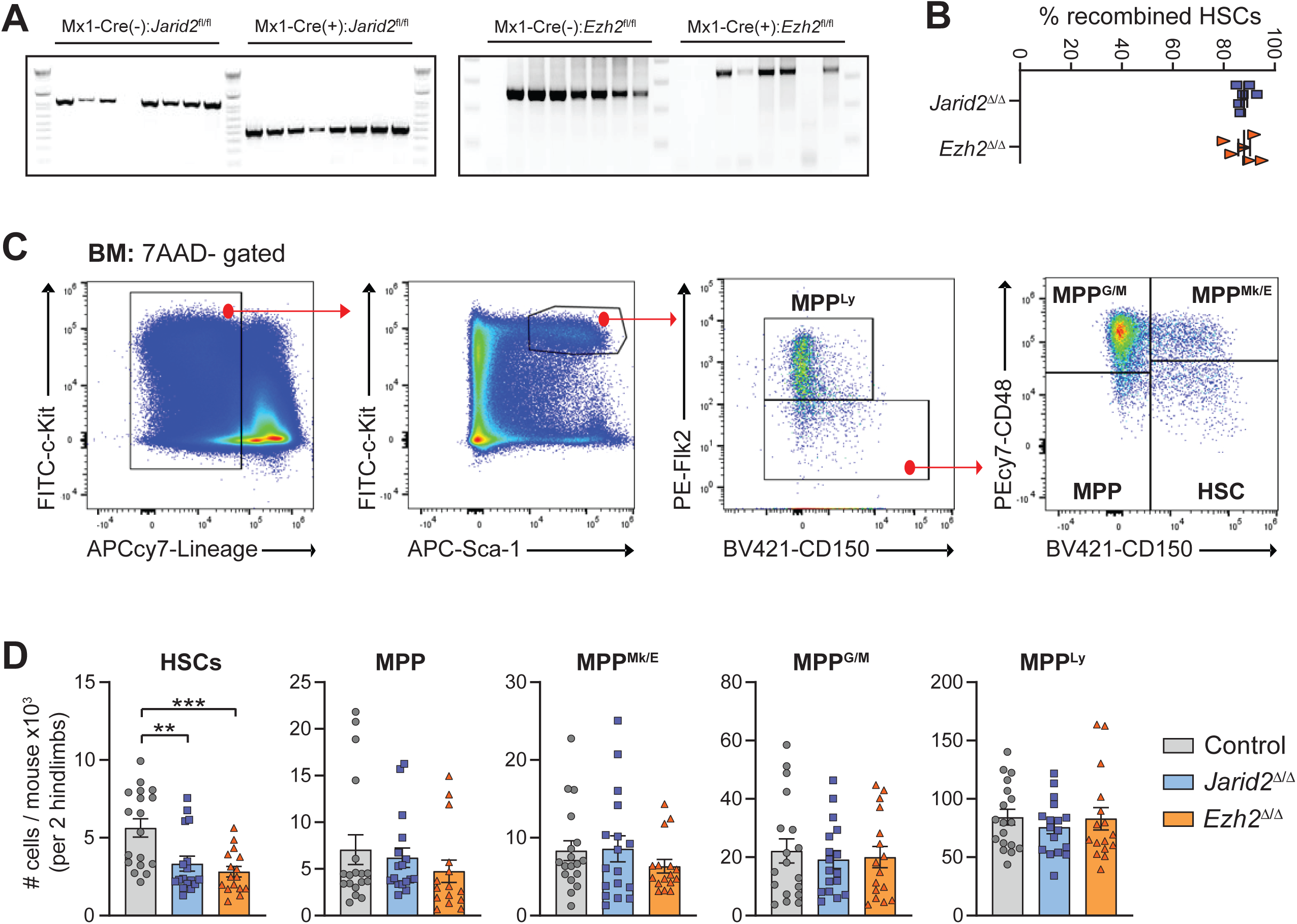
Genetic Deletion of Both *Jarid2* and *Ezh2* Lead to Depletion of Phenotypically-Defined HSCs. A) Genotyping PCR confirming successful excision of *Jarid2* (exon 4) and *Ezh2* (exons 16 and 17) in single HSC-derived colonies. B) Percentage of recombined single HSC-derived colonies per 96-well plate. Two biological replicates per three independent experiments. C) Representative flow cytometry gating strategy to identify indicated hematopoietic stem and progenitor cell populations. D) Absolute numbers of indicated HSPC populations from control (n=18), *Jarid2*^Δ/Δ^ (n=17) and *Ezh2*^Δ/Δ^ (n=16) mice eight-weeks post-pIpC. Data are mean ± s.e.m. * *p*<0.05, ** *p*<0.01, *** *p*<0.001, **** *p*<0.0001. One-way ANOVA with Tukey correction for multiple comparisons.

### Loss of *Ezh2* Does Not Convey Long-Term Reconstitution to MPPs Unlike Loss of Jarid2

A defining functional property resulting from conditional deletion of *Jarid2* in hematopoietic cells is the ability of MPP cells to engraft long-term in serial repopulation assays. To determine if this phenotype was related to PRC2, 100 MPPs (CD45.2+ Lineage-Sca-1+ c-Kit+ Flk2-CD48-CD150-) were isolated from Mx1-Cre control, *Jarid2*^Δ/Δ^ and *Ezh2*^Δ/Δ^ mice and competitively transplanted with 2.5×10^5^ congenic (CD45.1) bone marrow (BM) cells into lethally irradiated mice. Peripheral blood analysis revealed significantly reduced chimerism from *Ezh2*^Δ/Δ^ MPPs relative to control and *Jarid2*^Δ/Δ^ MPPs (**Fig. 2A**). But *Jarid2*^Δ/Δ^ MPPs were the only genotype capable of robust long-term multi-lineage reconstitution (**Fig. 2B,C**). Despite overall lower peripheral blood engraftment, *Ezh2*^Δ/Δ^ MPPs actually showed enhanced myeloid differentiation (**Fig. 2B**) and increased BM engraftment (**Fig. 2D**), which is composed mostly of mature myeloid cells. However, these cells could not be defined as functional repopulating cells as the lymphoid deficiency precluded long-term multi-lineage (LTMR; >1% donor-derived chimerism to myeloid, B-cell and T-cell lineages) reconstitution classification (**Fig. 2C**). Analysis 18-weeks post-transplant demonstrated increased numbers of donor-derived *Jarid2*^Δ/Δ^ and *Ezh2*^Δ/Δ^ MPPs in the BM of recipient mice (**Fig. 2E**). To determine if these *Ezh2*^Δ/Δ^ MPPs possessed self-renewal ability, 3.0×10^6^ BM cells were transferred from primary recipient to secondary irradiated mice. *Jarid2*^Δ/Δ^ MPPs were the only genotype to show robust peripheral blood engraftment in secondary recipients (**Fig. 2F**), with chimerism in all major blood lineages (**Fig. 2G**) and LTMR (**Fig. 2H**). While *Ezh2*^Δ/Δ^ MPPs showed some BM engraftment (**Fig. 2I**) and regeneration of donor-derived MPPs (**Fig. 2J**), the reduced LTMR strongly suggests that the function of Jarid2 in MPPs is largely PRC2-independent.

**Figure 2:**
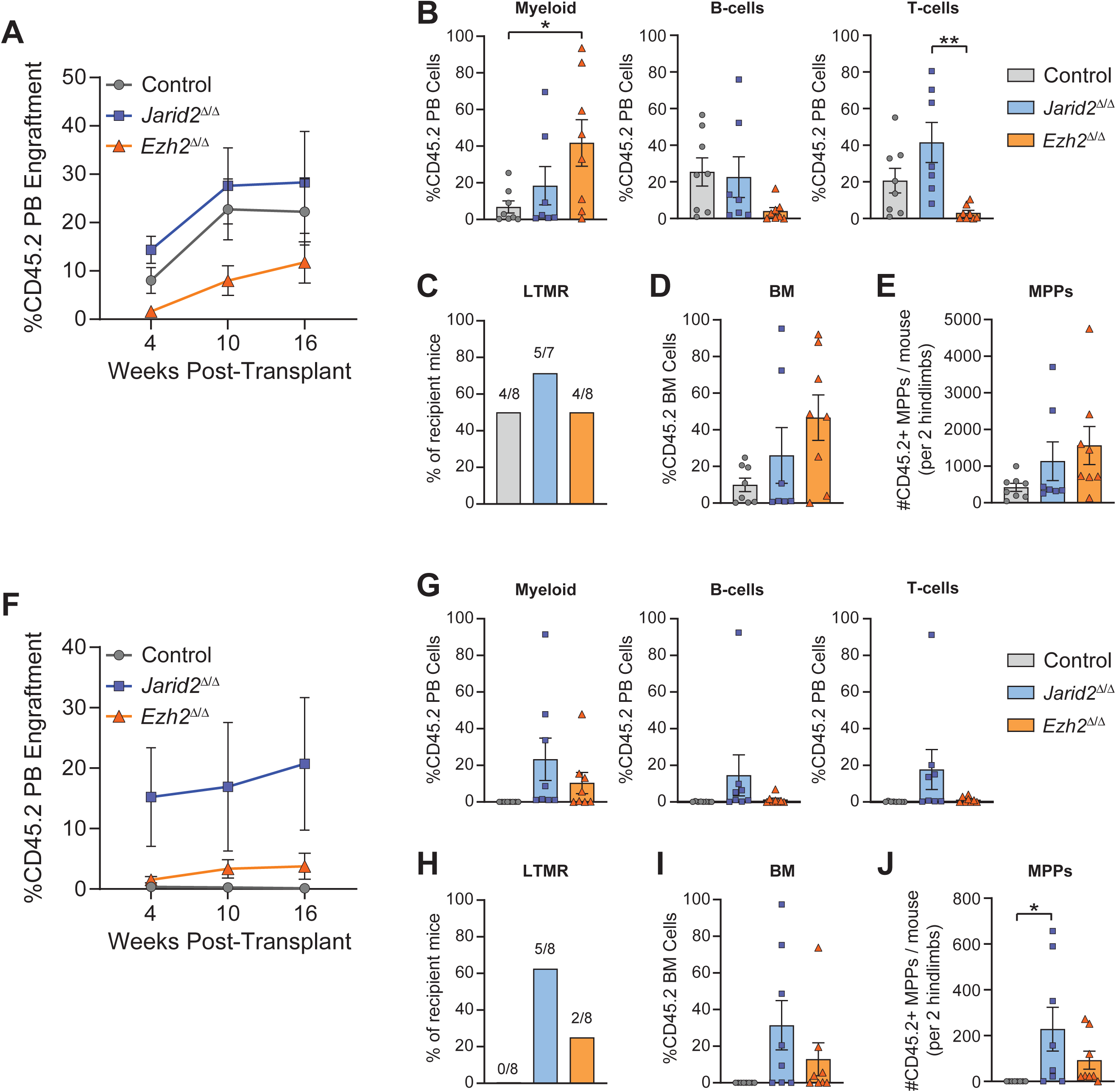
Loss of *Ezh2* Does Not Convey Long-Term Reconstitution to MPPs Unlike Loss of *Jarid2*. A) Percentage of donor-derived (CD45.2) peripheral blood (PB) engraftment from transplantation of 100 control (n=8), *Jarid2*^Δ/Δ^ (n=7) or *Ezh2*^Δ/Δ^ (n=8) MPPs in primary recipients. B) Donor-derived chimerism in myeloid, B-cell and T-cell PB lineages 16-weeks post-primary transplantation. C) Percentage of primary recipient mice with long term multi-lineage reconstitution (LTMR). D) Donor-derived bone marrow (BM) engraftment 18-weeks post-primary transplantation. E) Donor-derived MPP cell count from BM of primary recipient mice 18-weeks post-transplantation. F) Percentage of donor-derived PB engraftment from secondary transplantation of primary BM from control (n=8), *Jarid2*^Δ/Δ^ (n=8) or *Ezh2*^Δ/Δ^ (n=8) MPP-transplanted primary recipients. G) Donor-derived chimerism in myeloid, B-cell and T-cell PB lineages 16-weeks post-secondary transplantation. H) Percentage of secondary recipient mice with LTMR. I) Donor-derived BM engraftment 18-weeks post-secondary transplantation. J) Donor-derived MPP cell count from BM of secondary recipient mice 18-weeks post-transplantation. Data are mean ± s.e.m. * *p*<0.05, ** *p*<0.01, *** *p*<0.001, **** *p*<0.0001. One-way ANOVA with Tukey correction for multiple comparisons

### Ezh2 Restricts Myeloid Differentiation in HSCs Whereas Jarid2 Restricts T-cell Potential

To compare the functional effects of loss of *Jarid2* versus loss of *Ezh2* in an unbiased fashion, competitive transplantation of unfractionated whole BM was performed by transplanting 5.0×10^5^ BM cells from pIpC-treated Mx1-Cre control, *Jarid2*^Δ/Δ^ and *Ezh2*^Δ/Δ^ mice with 5.0×10^5^ BM cells from congenic mice. Loss of *Jarid2* enhanced peripheral blood engraftment (**Fig. 3A**) and chimerism in all major blood lineages (**Fig. 3B**), likely via the previously described increased function of *Jarid2*^Δ/Δ^ MPPs (**Fig. 2**). In contrast, *Ezh2* deficiency selectively increased myeloid differentiation without significantly altering lymphoid output compared to control BM (**Fig. 3B**). Both *Jarid2*^Δ/Δ^ and *Ezh2*^Δ/Δ^ genotypes showed significantly enhanced BM chimerism relative to control cells at 18-weeks post-transplant (**Fig. 3C**). While loss of *Ezh2* promoted general expansion of the collective hematopoietic stem and progenitor cells (HSPCs; Linage-Sca-1+ c-Kit+; **Fig. 3D**), *Ezh2*^Δ/Δ^ cells showed reduced donor-derived HSC regeneration in the BM of recipient mice (**Fig. 3E**) as we previously reported for *Jarid2*^Δ/Δ^ cells.

**Figure 3:**
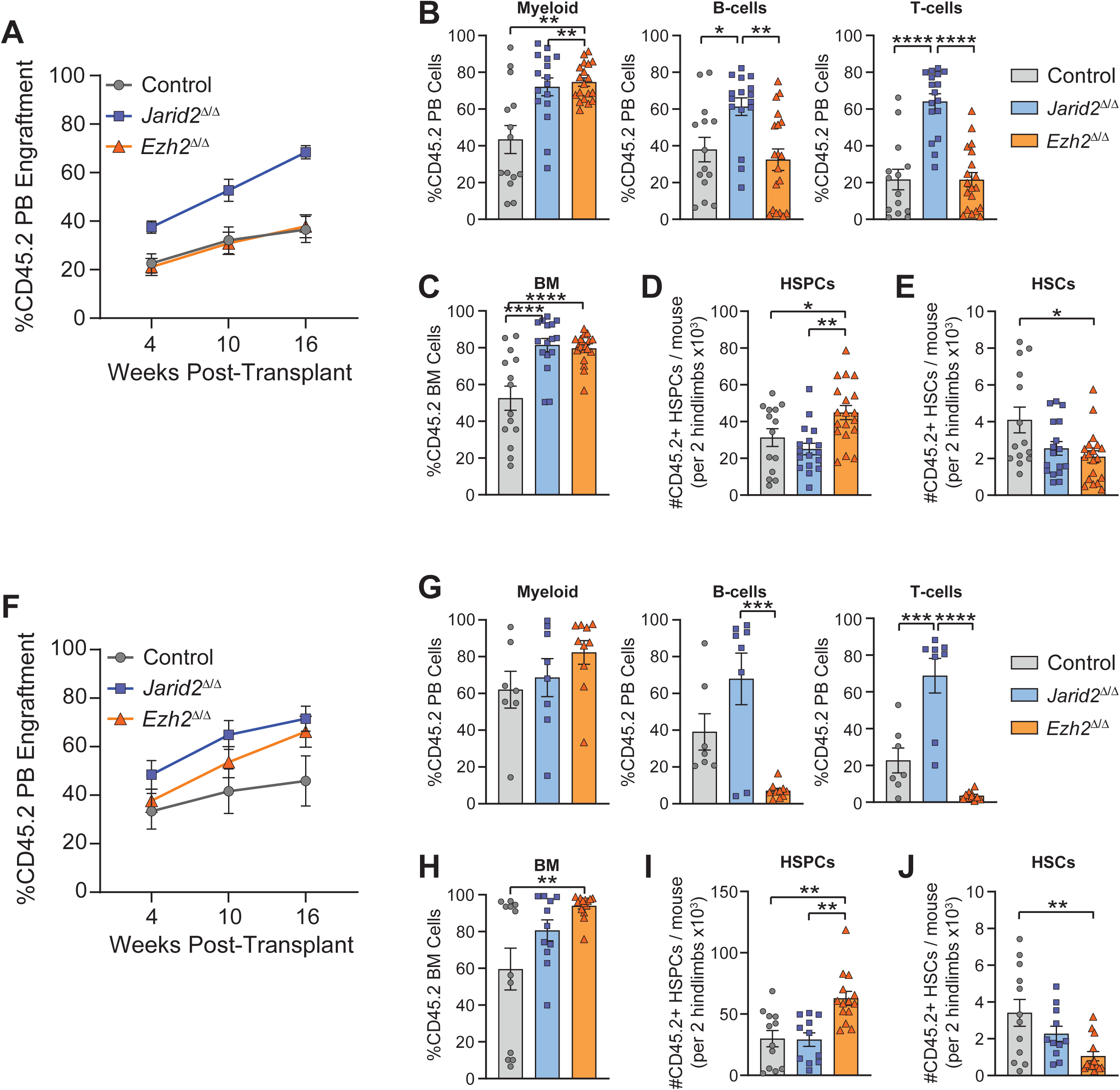
Ezh2 Restricts Myeloid Differentiation in HSCs Whereas Jarid2 Restricts T-cell Potential. A) Percentage of donor-derived (CD45.2) peripheral blood (PB) engraftment from competitive transplantation of WBM from control (n=14), *Jarid2*^Δ/Δ^ (n=16) or *Ezh2*^Δ/Δ^ (n=19) mice in primary recipients. B) Donor-derived chimerism in myeloid, B-cell and T-cell PB lineages 16-weeks post-primary transplantation. C) Donor-derived bone marrow (BM) engraftment 18-weeks post-primary transplantation. D) Donor-derived HSPC (CD45.2+ Lineage-Sca-1+ c-Kit+) cell count from BM of primary recipient mice 18-weeks post-transplantation. E) Donor-derived HSC (CD45.2+ Lineage-Sca-1+ c-Kit+ CD48-CD150+) cell count from BM of primary recipient mice 18-weeks post-transplantation. F) Percentage of donor-derived PB engraftment from secondary transplantation of primary BM from control (n=7), *Jarid2*^Δ/Δ^ (n=8) or *Ezh2*^Δ/Δ^ (n=10) WBM-transplanted primary recipients. G) Donor-derived chimerism in myeloid, B-cell and T-cell PB lineages 16-weeks post-secondary transplantation. H) Donor-derived BM engraftment 18-weeks post-secondary transplantation. I) Donor-derived HSPC cell count from BM of secondary recipient mice 18-weeks post-transplantation. J) Donor-derived HSC cell count from BM of secondary recipient mice 18-weeks post-transplantation. Data are mean ± s.e.m. * *p*<0.05, ** *p*<0.01, *** *p*<0.001, **** *p*<0.0001. One-way ANOVA with Tukey correction for multiple comparisons

To verify the observed phenotypes were heritable, secondary transplantation was performed by transferring 3.0×10^6^ unfractionated BM cells from individual primary recipients and transferring them each to unique secondary recipient mice. While overall peripheral blood engraftment was not significantly different (**Fig. 3F**), lineage bias was dramatically affected by loss of *Ezh2*. Chimerism of *Ezh2*^Δ/Δ^ cells was strongly biased towards myeloid differentiation with a severe deficiency in peripheral lymphoid output (**Fig. 3G**). In contrast, loss of *Jarid2* significantly enhanced T-cell potential (**Fig. 3G**). Due to the myeloid bias, *Ezh2*^Δ/Δ^ cells were dominant in the overall BM (**Fig. 3H**). HSPC expansion (**Fig. 3I**) and HSC depletion (**Fig. 3J**) of *Ezh2*^Δ/Δ^ cells was observed in an analogous fashion to primary transplantation. These results confirm the observed phenotypes are programmed at the level of long-term HSCs. Collectively, these data reveal a dichotomy where *Ezh2* (presumably PRC2-mediated) is required to restrict myeloid differentiation in HSCs, whereas the PRC2 co-factor Jarid2 normally restricts T-cell potential, potentially through a PRC2-independent mechanism.

### Loss of *Ezh2* Leads to Accumulation of Lymphoid-Biased MPP4 Cells

While loss of *Jarid2* increased overall differentiation output into all blood lineages, loss of *Ezh2* manifested a paradoxical reduction in long-term HSCs, but an expansion of total HSPCs which was associated with myeloid differentiation bias. To understand the mechanisms underlying this and ascertain which functions of Jarid2 may be PRC2-dependent in HSPCs, single cell transcriptomic analysis was performed on donor-derived HSPCs (CD45.2+ Lineage-Sca-1+ c-Kit+) obtained from primary recipients of competitive WBM transplantation (**Fig. 3**). Cell clustering and annotation were performed based on transcriptomic profiles, with dimensionality reduction visualized using uniform manifold approximation and projection (UMAP). Differential gene expression analysis between clusters was conducted using the Wilcoxon rank-sum test, applying thresholds of adjusted *p* < 0.01 and absolute log2 fold change > 0.25. Marker genes for each cluster were identified using the same criteria. Clusters were annotated by comparing enriched marker genes with established signatures of hematopoietic stem and progenitor cell (HSPC) populations (**Supplementary Table S1**)(Collins et al., 2024; Giladi et al., 2018). UMAP clustering confirmed the reduction in *Ezh2*^Δ/Δ^ HSCs observed by flow cytometry (**Fig. 4A**). However, there was a striking enrichment of *Ezh2*^Δ/Δ^ cells in the cluster annotated as MPP4 (**Fig. 4B**). This cell population, defined by expression of marker genes such as *Dntt*, *Flt3* and *Notch1* (**Fig. S1A**), are known to be lymphoid-biased MPPs (also called MPP^Ly^; **Fig. 1D**)(Challen et al., 2021; Pietras et al., 2015).

**Figure 4:**
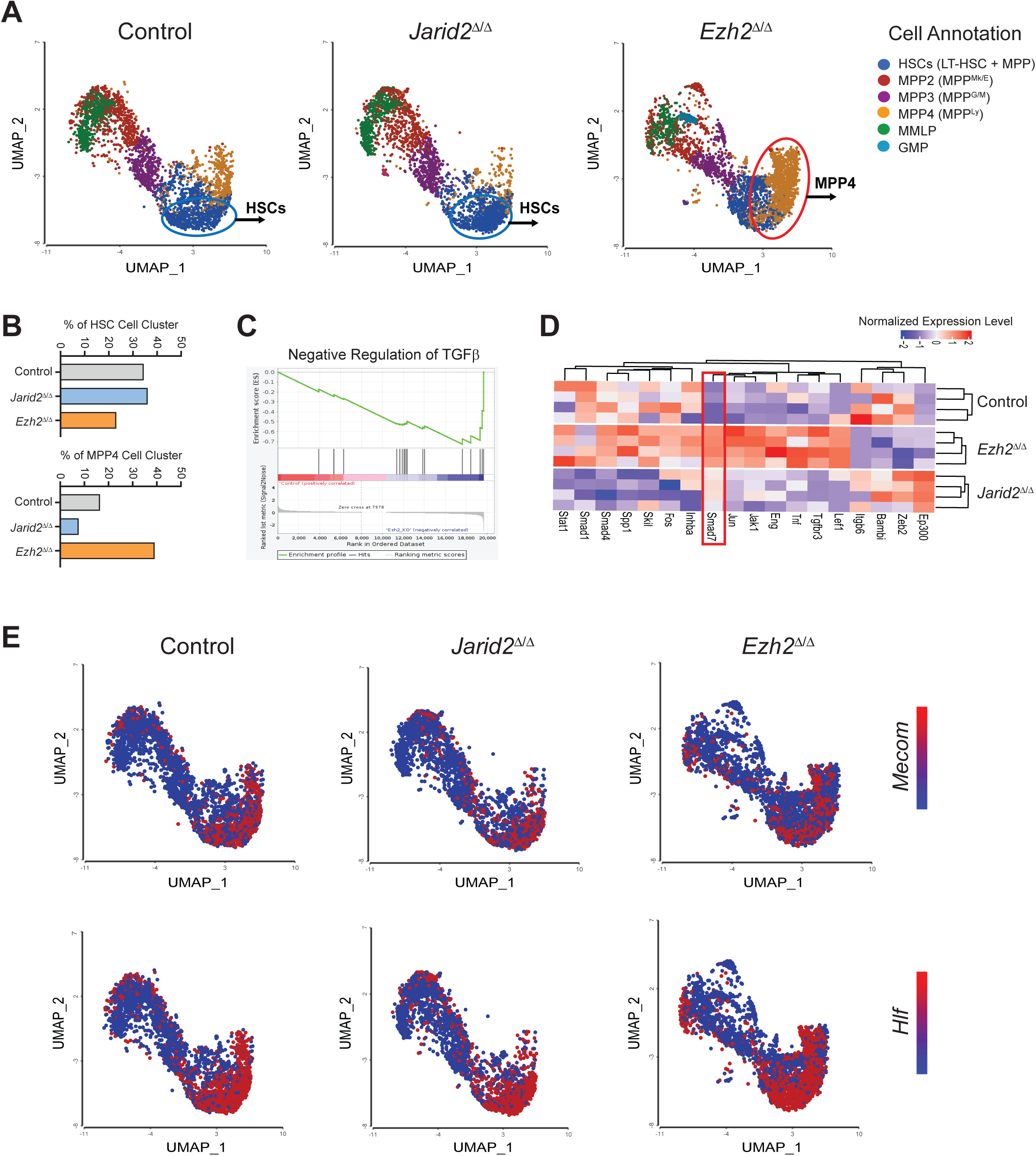
Loss of Ezh2 Leads to Accumulation of Lymphoid-Biased MPP4 Cells. A) UMAP clustering and cell annotation of scRNA-seq data generated from donor-derived HSPCs (CD45.2+ Lineage-Sca-1+ c-Kit+) post-primary transplant. B) Percentage cells of indicated genotypes within HSC and MPP4 cell clusters. C) Gene Set Enrichment Analysis (GSEA) of MPP4 cell cluster showing negative regulation of TGFβ signaling in *Ezh2*^Δ/Δ^ HSPCs compared to control cells. D) Pseudobulk differential gene expression analysis of MPP4 cluster cells showing upregulation of *Smad7* in *Ezh2*^Δ/Δ^ cells. E) Expression of *Mecom* and *Hlf* in cells of each genotype across the UMAP. Data are mean ± s.e.m. * *p*<0.05, ** *p*<0.01, *** *p*<0.001, **** *p*<0.0001. One-way ANOVA with Tukey correction for multiple comparisons

Transcriptomic comparison of the MPP4 genotypes by Gene Set Enrichment Analysis (GSEA) revealed the most significantly perturbed pathway to be downregulation of TGFβ signaling in *Ezh2*^Δ/Δ^ MPP4 cells (**Fig. 4C**). TGFβ signaling plays an essential role in regulating homeostasis of cells in the lymphoid lineage (Marie et al., 2005; Moreau et al., 2022). As an upstream signal, differential gene expression identified marked upregulation of *Smad7* in *Ezh2*^Δ/Δ^ MPP4 cells (**Fig. 4D**). Smad7 is an inhibitory Smad that suppresses TGFβ signaling by blocking Smad2/3 interaction with Smad4 and preventing downstream transcriptional activity (Hayashi et al., 1997; Itoh and ten Dijke, 2007). These findings suggest PRC2-mediated repression of *Smad7* is required to allow TGFβ-mediated lymphoid differentiation of MPP4 cells. Further analysis showed the accumulation of *Ezh2*^Δ/Δ^ MPP4 was associated with increased expression of genes typically associated HSC identity such as *Mecom*, *Hlf*, *Meis1* and *Hoxa9* (**Fig. 4E**; **Fig. S1B**), possibly implying that in the absence of *Ezh2*, these MPP4 acquire transcriptional features of HSCs which may convey some measure of self-renewal capacity. Taken together, these data indicate loss of *Ezh2* leads to differentiation block and accumulation of MPP4 cells due to de-repression of HSC-specific gene signature and failure of TGFβ-mediated lymphoid priming respectively.

### Loss of *Ezh2* Leads to Accumulation of B-cell Progenitors in the Bone Marrow

To further investigate the potential impacts of *Jarid2* and *Ezh2* loss-of-function on HSPC lineage priming in a native setting, Cellular Indexing of Transcriptomes and Epitopes by Sequencing (CITE-seq) was performed on HSPCs (Lineage-Sca-1+ c-Kit+) from sex-matched Mx1-Cre control, *Jarid2*^Δ/Δ^ and *Ezh2*^Δ/Δ^ mice to gain simultaneous information on gene expression and surface proteins at the single cell level. Antibody-derived tags were included for well-established cell surface markers of HSPC populations (c-Kit, Sca-1, CD48, CD150, CD201, and CD135; **Fig. S2A**). UMAP clustering and cell-type annotation were performed using both transcriptomic and surface protein (ADT) expression profiles. Cell clusters were annotated by integrating canonical gene expression patterns and HSPC surface protein markers (**Supplementary Table S2**)(Collins *et al*., 2024; Giladi *et al*., 2018).

As an initial quality control, potential sex-specific effects were examined. No significant differences were observed between males and females within each genotype (**Fig. 5A**). The most striking result was enrichment of two distinct B-cell progenitor cell clusters in *Ezh2*^Δ/Δ^ HSPCs (consistent across both sexes; **Fig. 5B, Fig. S2B**), defined by expression of B-lineage markers such as *Ebf1*, *Pax5* and *CD19* (**Fig. 5C**). Given the absence of sex-based bias, data from male and female mice were combined for each genotype in downstream analyses. GSEA demonstrated that *Ezh2* loss-of-function led to activation of B-cell survival pathways (**Fig. 5D**), driven by upregulation of critical survival genes such as *Mcl1* and *IL7r* (**Fig. 5E**). These data suggest loss of *Ezh2* promotes expansion of early B-cell progenitors in the BM through upregulation of factors that promote B-cell survival and proliferation.

**Figure 5:**
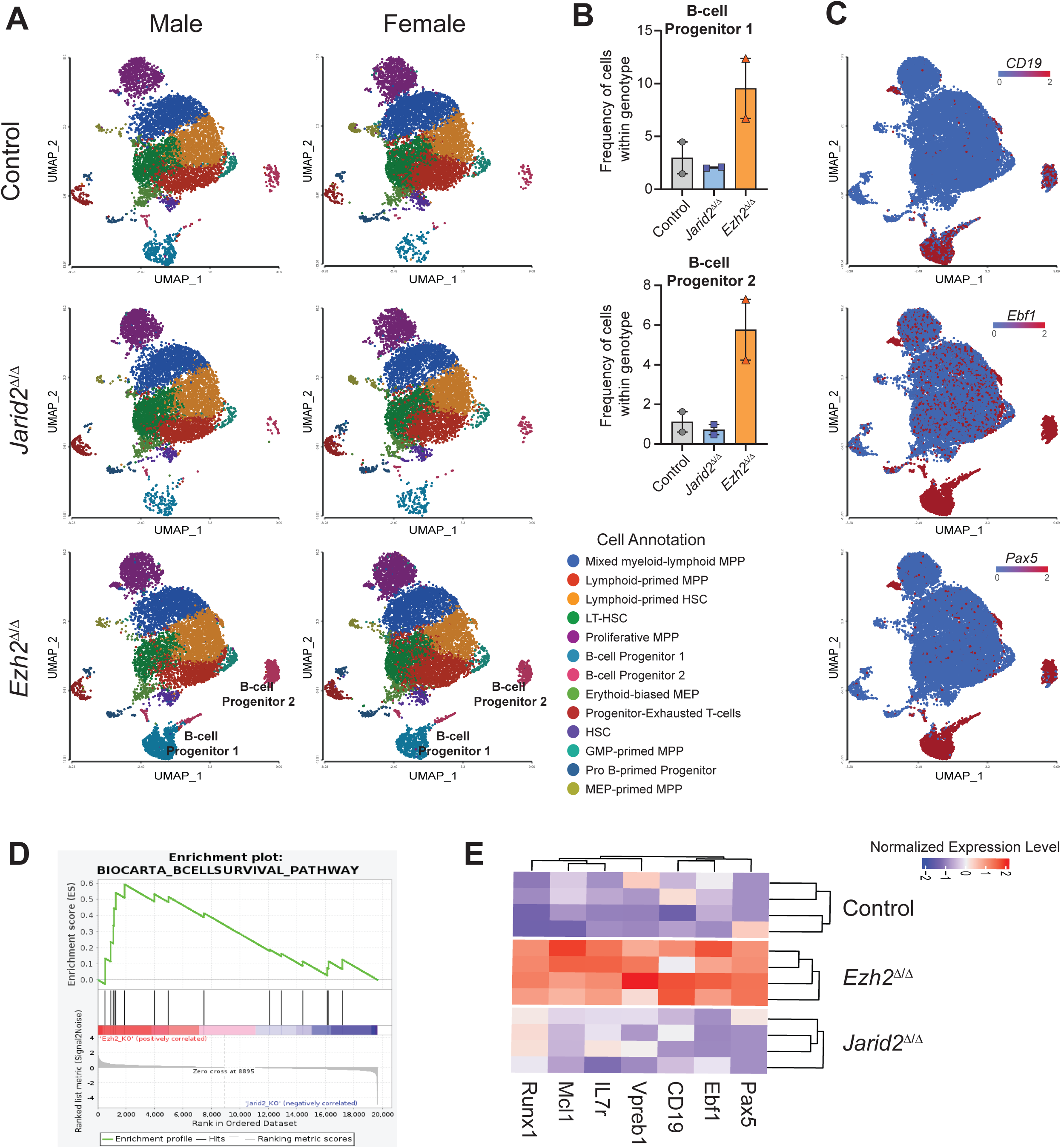
Loss of Ezh2 Leads to Accumulation of B-cell Progenitors in the Bone Marrow. A) UMAP clustering and cell annotation of CITE-seq data generated from HSPCs (Lineage-Sca-1+ c-Kit+) of indicated genotypes isolated from mice eight-weeks post-pIpC. B) Frequency of cell of indicated genotypes within B-cell Progenitor 1 and B-cell Progenitor 2 cell clusters. C) Expression of B-cell markers *CD19*, *Ebf1* and *Pax5* in cells across the UMAP. D) Gene Set Enrichment Analysis (GSEA) of B-cell Progenitor 1 and B-cell Progenitor 2 clusters showing enrichment of B-cell survival pathway gene expression in *Ezh2*^Δ/Δ^ cells. E) Pseudobulk differential gene expression analysis in B-cell Progenitor 1 and B-cell Progenitor 2 clusters showing overexpression of early B-cell survival genes in *Ezh2*^Δ/Δ^ cells. Data are mean ± s.e.m. * *p*<0.05, ** *p*<0.01, *** *p*<0.001, **** *p*<0.0001. One-way ANOVA with Tukey correction for multiple comparisons

### Loss of *Jarid2* Increases B-cell Output Whereas Loss of *Ezh2* Leads to Developmental Arrest of Pre-Pro B-cells

To determine if the transcriptional alterations in *Ezh2*^Δ/Δ^ HSPCs contributed to the functional deficiencies in lymphoid output observed from *Ezh2*^Δ/Δ^ cells in transplantation assays, B-cell developmental stages were examined in primary recipient mice by flow cytometry (**Fig. 6A**; **Fig. S3**). *Ezh2*^Δ/Δ^ recipient mice exhibited a significant accumulation of Pre-Pro B cells (B220+ IgM-CD43+ CD19-) compared to both control and *Jarid2*^Δ/Δ^ recipients (**Fig. 6B**). Further dissection of the Pre-Pro B-cell compartment by CD24 expression (Ayre et al., 2015) demonstrated *Ezh2* loss led to a marked increase in both early (CD24-) and late (CD24+) Pre-Pro B-cell populations (**Fig. 6B**). However, at later stages of B-cell development there was a significant reduction of *Ezh2*^Δ/Δ^ cells, in stark contrast to *Jarid2*^Δ/Δ^ cells which were significantly increased (**Fig. 6B**). To directly assess if loss of *Ezh2* induces maturational arrest of B-cell progenitors, *in vitro* B-cell differentiation assays were performed. Purified control, *Jarid2*^Δ/Δ^ and *Ezh2*^Δ/Δ^ HSCs were cultured on OP9 stromal cells with lymphoid cytokines and assessed for differentiation status by flow cytometry (**Fig. 6C**). After 10 days *in vitro*, there was a significant increase in early Pre-Pro B-cells in *Ezh2*^Δ/Δ^ co-cultures with a concomitant decrease in late Pre-Pro B-cells compared to control and *Jarid2*^Δ/Δ^ cells (**Fig. 6D**), directly confirming that *Ezh2* is required for appropriate B-cell maturation. In stark contrast, *Jarid2* appears to restrict B-cell potential in early HSPCs as loss of *Jarid2* consistently leads to increased B-cell differentiation output.

**Figure 6:**
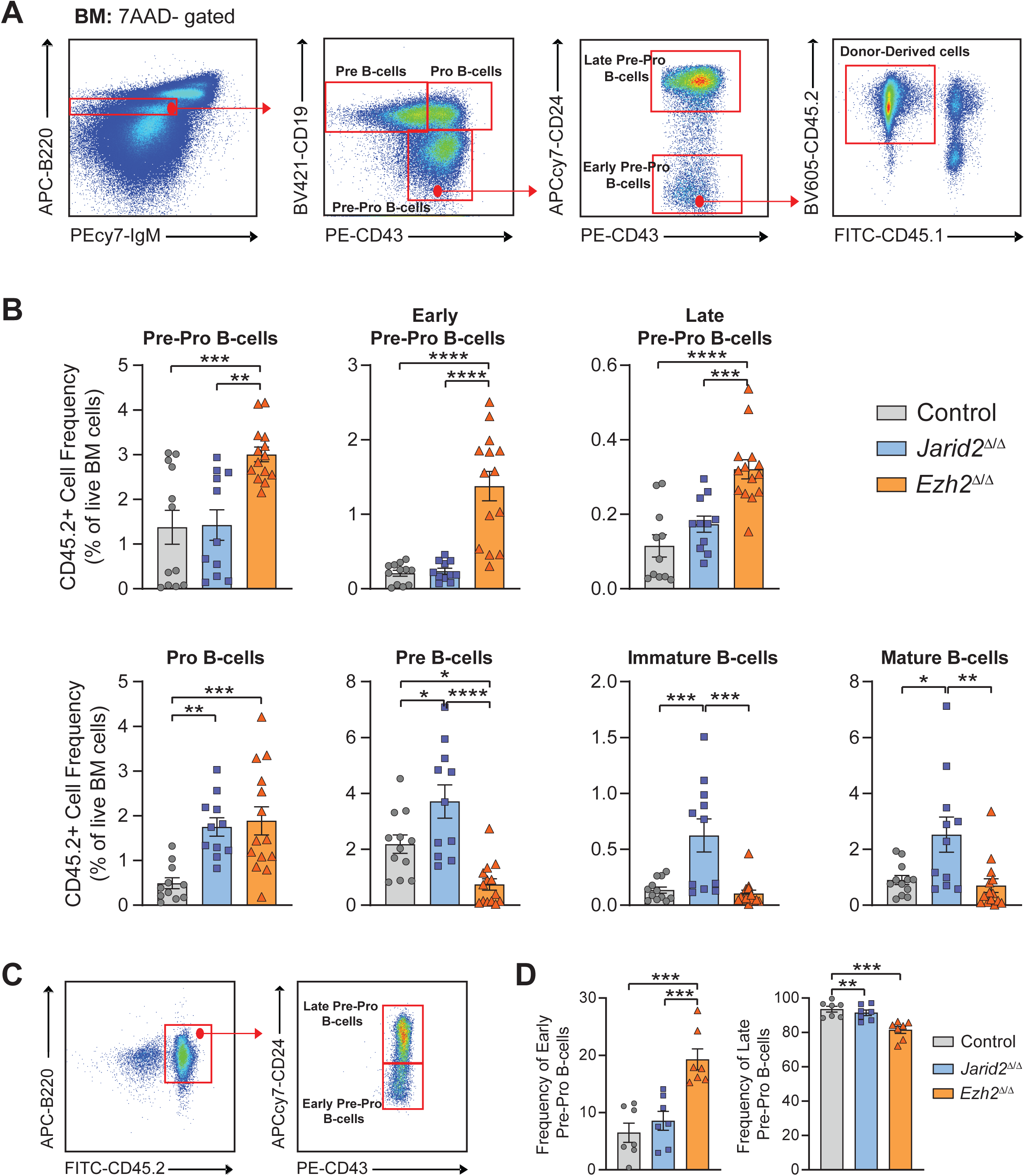
Loss of Jarid2 Increases B-cell Output Whereas Loss of Ezh2 Leads to Developmental Arrest of Pre-Pro B-cells. A) Representative flow cytometry gating to identify B-cell developmental stages in the bone marrow (BM) of recipient mice. B) Frequency of different donor-derived B-cell progenitor populations in the BM of primary recipient mice. C) Representative flow cytometry analysis of Day 10 cells from B-cell differentiation assays. D) Frequency of early and late Pre-Pro B-cells derived from HSCs of indicated genotypes at Day 10 of B-cell differentiation assay. Data are mean ± s.e.m. * *p*<0.05, ** *p*<0.01, *** *p*<0.001, **** *p*<0.0001. One-way ANOVA with Tukey correction for multiple comparisons

## DISCUSSION

Our prior work revealed the critical importance of the PRC2 co-factor Jarid2 in normal and neoplastic hematopoiesis. While *Jarid2* was demonstrated to be a *bona fide* tumor suppressor in chronic myeloid neoplasms, loss of *Jarid2* in normal MPPs leads to de-repression of HSC-specific transcriptional programs, conveying ectopic long-term self-renewal potential to this progenitor cell population (Celik *et al*., 2018). Our molecular analyses suggested only a fraction of the transcriptional changes arising from loss of *Jarid2* in MPPs were associated with changes in distribution of the repressive chromatin mark H3K27me3 mediated by PRC2. The goal of this study was to determine if Jarid2 may have PRC2-independent functions in hematopoiesis by comparing the functional and molecular properties of HSPCs lacking either *Jarid2* or *Ezh2*, the enzymatic component of PRC2. Our findings reveal strikingly divergent roles for *Ezh2* and *Jarid2* in regulating HSPC developmental maturation and lineage differentiation, nominating previously unrecognized PRC2-independent functions of Jarid2 in hematopoiesis.

The critical importance of PRC2-mediated gene regulation in hematopoiesis is underscored by the recurrence of *EZH2* somatic mutations in blood cancers. *EZH2* gain-of-function mutations are recurrent drivers of lymphoma (Beguelin et al., 2013; Morin *et al*., 2010) whereas EZH2 loss-of-function mechanisms (either point mutations or deletions of chromosome 7q) are common in myeloid neoplasms such as myelodysplastic syndromes (MDS) and myelofibrosis (MF) (Ernst et al., 2010; Nikoloski *et al*., 2010; Score *et al*., 2012). Despite this, there has been surprisingly little functional characterization of how *Ezh2* loss-of-function influences normal HSC fate decisions. A previous study suggested *Ezh2* loss-of-function did not dramatically alter function of HSCs in transplant assays (Xie *et al*., 2014). But these experiments were done with fetal liver or neonatal derived HSCs. This study also used *Ezh2*^fl/fl^ mice crossed to the Vav-Cre driver which we were unable to generate in this study, perhaps due to strain or animal house differences. Another study reported that over-expression of *Ezh2* by retroviral transduction increased transplantability of HSCs (Kamminga et al., 2006). We attempted to reconcile these differences and provide a definitive functional assessment of *Ezh2*-deficient HSPCs by using conditional knockout adult Mx1-Cre:*Ezh2*^fl/fl^ mice. Our findings demonstrate that *Ezh2* is necessary to restrict myeloid potential in HSCs. In the absence of *Ezh2*, HSC commitment becomes strongly biased towards myeloid differentiation and there is an extreme deficiency in lymphoid output in serial transplant. This deficit in peripheral lymphoid cells was underpinned by a developmental arrest of lymphoid-biased MPP4 and Pre-Pro B-cells in the absence of *Ezh2*. At the molecular level, this was associated with repression of TGFβ signaling and increased expression of B-cell survival and proliferation gene expression programs. The phenotypes are in stark contrast to HSPCs conditionally deficient for the PRC2 co-factor *Jarid2*. In the absence of *Jarid2*, differentiation into all peripheral blood lineages is enhanced with a particular bias towards T-cell generation. These contrasting functional phenotypes strongly suggest that Jarid2 may regulate HSC fate decisions at least partially via PRC2-independent mechanisms. However, both loss of *Jarid2* and *Ezh2* led to a reduction of phenotypically-defined HSCs, although this did not appear to compromise overall blood regeneration. We attempted to reconcile the identity of this depleted HSC subpopulation with CITE-seq profiling but were unable to define a connection. This may suggest that some aspect of HSC pool maintenance is mediated by PRC2-dependent Jarid2 functions. Further work is needed to elucidate the identity of this HSC subpopulation which may be important under various environmental stresses beyond the scope of the current work.

This work provides clear resolution into the function of *Ezh2* in adult HSCs during normal hematopoiesis and provides evidence supporting the concept of non-canonical functions of Jarid2 regulating HSC fate decisions. This adds to the growing body of literature suggesting many epigenetic regulators that are molecularly defined in biochemical or isolated cell culture assays may have functions outside these described roles that regulate stem cell populations *in vivo*. Future studies are warranted to dissect the chromatin-level mechanisms underlying the observed differences and clarify how Ezh2 facilitate B-lineage priming in HSPCs. Together, this work positions *Jarid2* and *Ezh2* as critical, mechanistically distinct regulators of HSC fate with broad relevance to the regulation of normal and neoplastic hematopoiesis.

## ACKNOWLEDGEMENTS

We thank all members of the Challen laboratory, particularly Samantha Burkart for laboratory management. We thank the Alvin J. Siteman Cancer Center, supported in part by an NCI Cancer Center Support Grant P30CA091842. This publication is solely the responsibility of the authors and does not necessarily represent the official views of the NIH. G.A.C was supported by the National Institutes of Health (NIH; HL147978, CA236819 and DK124883), the Edward P. Evans Foundation, the Leukemia and Lymphoma Society (6667-23) and the American Cancer Society (CSCC-RSG-23-991417-01-CSCC). H.B. was supported by the Edward P. Evans Center for MDS at Washington University in St. Louis. I.X.R was supported by NIH P30 CA091842. A.K. was supported by the American Society of Hematology (Graduate Hematology Award). J.A.M. was supported by the NIH (HL152180) and the Children’s Discovery Institute of Washington University and St. Louis Children’s Hospital. J.A.M. is a Scholar of the Leukemia and Lymphoma Society.

## AUTHOR CONTRIBUTIONS

Conceptualization and study design – G.A.C.

Experimentation and data acquisition – H.B., W.H., N.I., A.K., I.X.R., J.A., Y.L.

Data analysis – H.B., W.H., W.Y., J.A.M., G.A.C.

Funding acquisition – G.A.C.

Project administration and supervision – G.A.C.

Manuscript preparation – H.B., G.A.C.

## DECLARATION OF INTERESTS

The authors declare the following competing interests (unrelated to this work): G.A.C. has performed consulting and received research funding from Incyte, Ajax Therapeutics, Atavistik Bio and ReNAgade Therapeutics Management, and is a co-founder, member of the scientific advisory board and shareholder of Pairidex, Inc.

## SUPPLEMENTARY TABLES

**Supplementary Table S1: *Differentially expressed genes across single-cell RNA sequencing cell clusters*.**

The table includes gene symbols, fold change, and associated p-values. These genes were used to annotate the six distinct clusters, with top marker genes selected based on previously published single-cell RNA-seq datasets. Cluster-specific gene markers were identified using the Wilcoxon rank-sum test, applying a significance threshold of adjusted *p* < 0.01 and an absolute log2 fold change >0.25.

**Supplementary Table 2. *Differentially expressed genes and antibody-derived protein markers across single-cell CITE-seq cell clusters***.

The table includes gene symbols, fold change, and associated p-values for both RNA and antibody capture features. These data were used to annotate the 13 clusters, with top gene and protein markers selected based on prior single-cell CITE-seq studies. Cluster-specific markers were identified using the Wilcoxon rank-sum test, with a significance threshold of adjusted *p* < 0.01 and an absolute log2 fold change >0.25.

## METHODS

### Animals

All animal procedures were approved by the Institutional Animal Care and Use Committee of Washington University. All mice used in this study were C57Bl/6 background. *Jarid2*^fl/fl^ mice were previously described (Mysliwiec et al., 2006). *Ezh2^fl/fl^* mice (Neff et al., 2012) were obtained from Dr. Jeffrey Magee (Washington University). *Jarid2* and *Ezh2* carrying homozygous floxed alleles were crossed to the Mx1-Cre driver. For strains carrying Mx1-Cre, recombination of floxed alleles (=“Δ”) was induced by intraperitoneal injection of six doses (300μg/mouse) of polyinosinic–polycytidylic acid (pIpC; Sigma #P1350) given every other day. 10-12 week old mice were typically used for experimentation. Equal numbers of male and female mice were used, no gender biases were noted. Eight-week old male and female C57Bl/6 CD45.1 mice (The Jackson Laboratory # 002014) were used as recipients for bone marrow transplantation assays.

### Methocult-based HSC Genotyping

Mx1-Cre induced recombination efficiency was assessed by sorting single HSCs from either *Jarid2*^Δ/Δ^ or *Ezh2*^Δ/Δ^ mice into 96-well plates containing Methocult M3434 medium (Stem Cell Technologies #03434). After two-weeks culture, individual colonies were collected and washed with Dulbecco’s Phosphate Buffer Saline (Sigma #D8537), and genomic DNA was isolated using the KAPA Express Extract Kit (Sigma # KK7103). To confirm the deletion of *Jarid2* (exon 3) and *Ezh2* (exons 16 and 17), PCR was performed as previously described using primer pairs: Jarid2-FloxOut-F//Jarid2-FloxOut-R (Mysliwiec et al., 2006) and Ezh2 Excision F1//Ezh2 Excision R1//Ezh2 Excision R2 (Neff et al., PNAS, 2012), respectively. The recombination efficiency was checked using the following PCR conditions: 95 °C for 3 mins, 95 °C for 15 sec, 60 °C for 15 sec, 72 °C for 20 sec, and 72 °C for 5 mins.

### Bone Marrow Transplantation

CD45.1 congenic C57Bl/6 mice were used as recipients for all transplantation experiments. Mice received a lethal dose of total body irradiation administered in two split doses of 5.25 Gy (10.5 Gy total) delivered ∼4 hours apart using a gamma irradiator. For primary competitive transplantation experiments, 100 donor-derived multipotent progenitor (MPP) cells (CD45.2^+^ Lin^−^ Sca-1^+^ c-Kit^+^ CD48^−^ CD150^−^) were isolated via fluorescence-activated cell sorting (FACS) on a MoFlo cell sorter (Beckman Coulter) and transplanted via retro-orbital injection along with 2.5×10⁵ unfractionated wild-type whole bone marrow (WBM) competitor cells. For primary whole bone marrow transplants, 5×10⁵ total CD45.2 WBM cells were co-transplanted with 5×10⁵ wild-type CD45.1 WBM competitor cells into lethally irradiated recipients. Donor and competitor cells were distinguished by differential expression of CD45 allelic isoforms: CD45.2 (donor) and CD45.1 (competitor and recipient). At 18 weeks post-transplant, 3×10⁶ WBM cells were harvested from individual primary recipients transferred into uniquely identified lethally irradiated secondary recipients to assess long-term repopulating capacity.

### Flow Cytometry

Peripheral blood (PB) was collected from recipient mice and red blood cells were lysed using ACK lysis buffer (Gibco). Leukocytes were stained for donor chimerism and lineage analysis using flow cytometry (Cytek Northern Lights). Surface markers used to assess lineage contribution included myeloid lineage markers Gr-1 (Ly6G/Ly6C) and Mac-1 (CD11b), B-cell lineage marker B220 (CD45R), and T-cell lineage marker CD3ε. CD45.1 and CD45.2 were used to discriminate recipient/competitor and donor-derived populations, respectively. At 18 weeks post-transplantation, recipient mice were sacrificed, and whole bone marrow (WBM) was harvested from femurs, tibias, and iliac crests. BM cells were stained for hematopoietic stem and progenitor cells (HSPCs) and mature lineage markers. Lineage markers included Gr-1, Mac-1, Ter119, CD3ε, B220. HSPC panel included c-Kit (CD117), Sca-1 (Ly6A/E), Flk2 (Flt3), CD150 (Slamf1), CD48 (SLAMF2). LSK (Lin^−^ Sca-1^+^ c-Kit^+^) populations and HSPC subfractions including MPP^Ly^ (Lin^−^ Sca-1^+^ c-Kit^+^ Flk2^+^ CD48^+^ CD150*^-^*), MPP^G/M^ (Lin^−^ Sca-1^+^ c-Kit^+^ Flk2^-^ CD48^+^ CD150*^-^*), MPP^Mk/E^ (Lin^−^ Sca-1^+^ c-Kit^+^ Flk2^-^ CD48^+^ CD150*^+^*), MPP (Lin^−^ Sca-1^+^ c-Kit^+^ Flk2^-^ CD48^-^ CD150*^-^*), and HSC (Lin^−^ Sca-1^+^ c-Kit^+^ Flk2^-^ CD48^-^ CD150*^+^*) were identified and quantified. Bone marrow B cell development was analyzed using B220, CD19, CD43, CD24, IgM, and IgD to identify pre-pro-B, early pre-pro-B, late pre-pro-B, pro-B, pre-B, immature, and mature B cell subsets. The standard Hardy fraction flow cytometry gating strategy was used to classify B cell developmental stages in the bone marrow.

All antibody staining was performed in HBSS buffer (Corning #21021CV) containing Pen/Strep (100 Units/mL; Fisher Scientific #MT30002CI), HEPES (10uM; Life Technologies # 15630080) and FBS (2%; Sigma #14009C). Briefly, cells were suspended in complete HBSS at a concentration of 5×10^8^ cells/mL and incubated on ice for 20 minutes with the desired antibodies listed in the table below. For cell sorting, magnetic enrichment was carried out using the AutoMACS Pro Seperator (Miltenyi Biotec) with mouse CD117-conjugated microbeads (Miltenyi Biotec #130-091-224). Post-enrichment, the positive cell fraction was stained with appropriate antibodies and sorted by FACS (Beckman Coulter MoFlo). All antibodies utilized in this study were used at 1:100 dilutions and were obtained from BioLegend, eBioscience or BD Biosciences unless otherwise stated. Samples were analyzed using a Cytek Northern Lights flow cytometer. Data were analyzed using FlowJo software (BD Biosciences). Fluorescence minus one (FMO) controls and single-color controls were included for compensation and gating.

| Flow Cytometry Antibody | Source | Identifier |
| --- | --- | --- |
| BV605 anti-mouse CD45.2 | Biolegend | Cat# 109841 |
| FITC anti-mouse CD45.1 | Biolegend | Cat# 110706 |
| APCy7 anti-mouse Gr-1 | Biolegend | Cat# 108424 |
| APCy7 anti-mouse Mac-1 | Biolegend | Cat# 101226 |
| APCy7 anti-mouse B220 | Biolegend | Cat# 103224 |
| APCy7 anti-mouse CD3e | Biolegend | Cat# 100330 |
| APCy7 anti-mouse Ter119 | Biolegend | Cat# 116223 |
| BV421 anti-mouse c-Kit | Biolegend | Cat# 105827 |
| APC anti-mouse Sca-1 | Biolegend | Cat# 122512 |
| PECy7 anti-mouse CD48 | Biolegend | Cat# 103424 |
| PE anti-mouse CD150 | Biolegend | Cat# 115904 |
| BV421 anti-mouse CD45.2 | Biolegend | Cat# 84208 |
| APC anti-mouse CD3e | Biolegend | Cat# 100312 |
| APC anti-mouse B220 | Biolegend | Cat# 103212 |
| PECy7 anti-mouse Gr-1 | Biolegend | Cat# 108416 |
| PECy7 anti-mouse Mac-1 | Biolegend | Cat# 101216 |
| PECy7 anti-mouse B220 | Biolegend | Cat# 103222 |
| PE anti-mouse Flk2 | BD Biosciences | Cat# 553842 |
| FITC anti-mouse c-Kit | BD Biosciences | Cat# 561680 |
| BV421 anti-mouse CD150 | Biolegend | Cat# 115925 |
| PECy7 anti-mouse IgM | eBioscience | Cat# 25-5790-82 |
| BV421 anti-mouse CD19 | Biolegend | Cat# 115538 |
| PE anti-mouse CD43 | eBioscience | Cat# 12-0431-83 |
| APCcy7 anti-mouse CD24 | Biolegend | Cat# 101850 |
| FITC anti-mouse CD45.2 | Biolegend | Cat# 109806 |
| PE anti-mouse IgD | eBioscience | Cat# 12-5993-82 |
| Biotin anti-mouse Sca-1 | Biolegend | Cat# 108103 |

### Single-cell RNA Sequencing (scRNA-seq)

c-kit+ cells were enriched from the bone marrow of 18-week post-primary transplant recipient mice. Donor-derived (CD45.2+) KSL (c-Kit+ Sca1+ Lineage–) cells were subsequently sorted from the enriched population and resuspended at a concentration of 1,200 cells/μL in PBS supplemented with 0.04% BSA. Single-cell libraries were prepared using the Chromium Next GEM Single Cell 3′ Reagent Kits v3.1 (Dual Index)(10x Genomics, PN-1000268), following the manufacturer’s protocol (User Guide CG000317 Rev D) with minor modifications. Cell partitioning and barcoding were performed using the Chromium Next GEM Chip G Single Cell Kit (PN-1000127) and Chromium Dual Index Kit TT Set A (PN-1000215). GEM generation and reverse transcription (GEM-RT), followed by post-GEM cleanup, were conducted according to standard procedures. Purified cDNA was amplified for 11–13 cycles based on input cell number and quality, and then cleaned using SPRIselect beads (Beckman Coulter, Cat. No. B23318). Amplified cDNA was assessed using the Agilent Bioanalyzer High Sensitivity DNA Kit (Agilent, Cat. No. 5067-4626) to determine concentration and size distribution. Final gene expression (GEX) libraries were constructed as per the 10x Genomics protocol, adjusting the number of amplification cycles according to cDNA input. The libraries were quantified using the KAPA Library Quantification Kit for Illumina platforms (Roche/KAPA Biosystems, Cat. No. KK4824) and sequenced on the Illumina NovaSeq X Plus system using the XP workflow on an S4 Flow Cell with a paired-end 28 × 10 × 10 × 150 bp sequencing strategy. A target median depth of 50,000 reads per cell was achieved.

FastQ files were demultiplexed, aligned to the mouse reference genome (mm10), and processed using Cell Ranger v3.1.0. After generation of the gene-barcode matrix, low-quality cells were filtered out based on quality control metrics: cells with >10% mitochondrial gene expression (indicative of stress or apoptosis) and cells in the top 5% of total UMI or feature counts (to exclude potential multiplets or doublets) were excluded from downstream analysis. Further analysis was performed using Seurat v3.0 in R and Partek Flow® software. Dimensionality reduction was conducted via principal component analysis (PCA), and clusters were visualized using both t-distributed stochastic neighbor embedding (t-SNE) and Uniform Manifold Approximation and Projection (UMAP). Marker genes for each cluster were identified using a Wilcoxon rank-sum test, with significance set at adjusted *p* < 0.01 and absolute log2 fold change >0.25. Clusters were annotated based on known hematopoietic stem and progenitor cell (HSPC) gene markers. Differential gene expression analysis within clusters was performed to compare cells from control, *Jarid2*^Δ/Δ^, and *Ezh2*^Δ/Δ^ mice. Enriched pathways were identified using the Gene Set Enrichment Analysis (GSEA) software package. All raw read data (FASTQ files) are publicly available at the Gene Expression Omnibus database under accession number GSE302605.

### Single-cell CITE-seq

c-Kit+ Sca1+ Lineage– (KSL) cells were sorted by FACS. Cells were first stained with biotin-conjugated anti-Sca1 (Ly-6A/E) antibody for 20 minutes at 4 °C, then washed and incubated with a panel of six TotalSeq™-B barcoded antibodies targeting surface markers: streptavidin-PE (Sca-1; BioLegend #405287), CD117 (BioLegend #105849), CD48 (BioLegend #103457), CD150 (BioLegend #115951), CD135 (BioLegend #135319), and CD201 (BioLegend #141511). Antibody staining was performed using 1 µL of each antibody in 50 µL of PBS containing 0.04% BSA, for 30 minutes on ice, protected from light. Following staining, cells were washed with cold PBS containing 0.04% BSA, filtered through a 40 µm cell strainer and assessed for viability (>90%) using Trypan Blue exclusion. After that, cells were resuspended at 1,200 cells/μL in PBS + 0.04% BSA. Single-cell transcriptome and protein profiling were performed using the Chromium Next GEM Single Cell 3′ Reagent Kits v3.1 (Dual Index) (PN-1000268) in conjunction with the Feature Barcode Kit for Cell Surface Protein (PN-1000262), following the manufacturer’s protocol (User Guide CG000317 Rev D) with slight optimizations to the staining and preparation steps. Cells were loaded onto the Chromium Controller using the Chromium Next GEM Chip G (PN-1000127), targeting recovery of approximately 10,000 cells per channel. Gel bead-in-emulsion (GEM) generation, barcoding, and reverse transcription were carried out using the recommended thermal cycling conditions: Lid temperature 53 °C, 53 °C for 45 min, and 85 °C for 5 min; hold at 4 °C. Post-RT GEMs were broken, and barcoded cDNA was purified using Dynabeads™ MyOne™ Silane (PN-2000048). Amplified cDNA containing both mRNA– and antibody-derived tags (ADTs) was generated via PCR using cycling conditions recommended by 10x Genomics and cleaned up using SPRIselect reagent (Beckman Coulter, Cat. No. B23318).

Separate libraries were prepared for gene expression and Feature Barcode (protein) profiling: 1. Gene Expression Library: A portion of the cDNA was enzymatically fragmented, end-repaired, A-tailed, and ligated to adapters, followed by sample index PCR amplification using the Dual index Kit TT Set A (PN-1000215). 2. Feature Barcode Library: A separate cDNA aliquot was amplified using primers targeting the Feature Barcode constructs using Dual index Kit NT Set A (PN-1000242). Final libraries were quantified using a Qubit 4 Fluorometer and assessed for size distribution using the Agilent Bioanalyzer High Sensitivity DNA Kit (Cat. No. 5067-4626). Libraries were pooled in appropriate ratios and sequenced on the Illumina NovaSeq X Plus system using an S4 flow cell with the following read configuration: 28 bp (Read 1), 10 bp (i7 index), 10 bp (i5 index), and 90 bp (Read 2). A target median sequencing depth of ∼50,000 reads per cell was used for gene expression and 5,000–10,000 reads per cell for antibody-derived tags.

FastQ files were demultiplexed and processed using Cell Ranger v3.1.0. Gene expression and antibody capture reads were analyzed simultaneously using the cellranger count pipeline with a custom reference genome built from the mm10 genome assembly and the appropriate Feature Barcode reference file. Cells with >10% mitochondrial transcript content and those within the top 5% of UMI counts (potential doublets) were excluded from downstream analysis.

Further analysis was performed using Seurat v3.0 (R package) and Partek Flow® software. Dimensionality reduction was conducted using principal component analysis (PCA) on the gene expression matrix, and antibody-derived tag (ADT) data were analyzed in parallel following centered log-ratio (CLR) normalization. Clustering was performed using graph-based algorithms and visualized via t-SNE and UMAP. Cluster-specific gene and surface protein markers were identified using the Wilcoxon rank-sum test, with significance thresholds of adjusted *p* < 0.01 and absolute log2 fold change > 0.25. Cluster annotation was guided by known hematopoietic stem and progenitor cell (HSPC) transcriptomic and surface marker profiles. Differential gene expression analysis within clusters was performed to compare cells from control, *Jarid2*^Δ/Δ^, and *Ezh2*^Δ/Δ^ mice. Enriched pathways were identified using the Gene Set Enrichment Analysis (GSEA) software package. All raw read data (FASTQ files) are publicly available at the Gene Expression Omnibus database under accession number GSE302605.

### OP9-Based B-Cell Differentiation Assay

To assess B-cell differentiation potential, an OP9 stromal cell co-culture system was employed. On Day 1, OP9 cells were seeded at a density of 5×10⁴ cells per well in 24-well plates using OP9 culture medium composed of α-MEM (Gibco), supplemented with 20% fetal bovine serum (FBS; Gibco) and 1% penicillin/streptomycin (Gibco). On Day 2, the medium was replaced with B-cell differentiation medium consisting of OP9 culture medium supplemented with 5 ng/mL Flt3 ligand (Flt3L; PeproTech) and 5 ng/mL interleukin-7 (IL-7; PeproTech). Immediately following the medium change, 250 HSCs were plated per well onto the OP9 monolayer. Cultures were maintained for 10 days, with B-cell differentiation medium replenished every 3 days. On Day 10, non-adherent and loosely adherent cells were harvested and filtered through a 40μm cell strainer (Corning) to remove residual OP9 stromal cells. Single-cell suspensions were stained with fluorophore-conjugated antibodies against CD45.2 and B-cell progenitor markers (B220, CD43, and CD24) for flow cytometric analysis.

### Quantification, UMAP Cell Clustering, and Statistical Analysis

Statistical comparisons were performed using one-way ANOVA followed by Tukey’s post hoc test for multiple comparisons. Cell clustering and annotation from single-cell RNA sequencing (scRNA-seq) and CITE-seq data were performed based on integrated gene expression and antibody-derived tag (ADT) profiles. Cluster-specific marker identification used the Wilcoxon rank-sum test with thresholds of adjusted *p* < 0.01 and absolute log2 fold change > 0.25. Pseudobulk differential expression analysis and gene set enrichment analysis (GSEA) were conducted using thresholds of adjusted *p* < 0.05 and absolute log2 fold change >0.25. Statistical significance is represented as follows: ∗*p*<0.05, ∗∗*p*<0.01, ∗∗∗*p*<0.001, ∗∗∗∗*p*<0.0001. Error bars on graphs represent the standard error of the mean (S.E.M.).

