## Supplementary Figures for "Functional Comparison to Ezh2 Reveals PRC2-Independent Functions of Jarid2 in Hematopoietic Stem Cell Lineage Commitment"

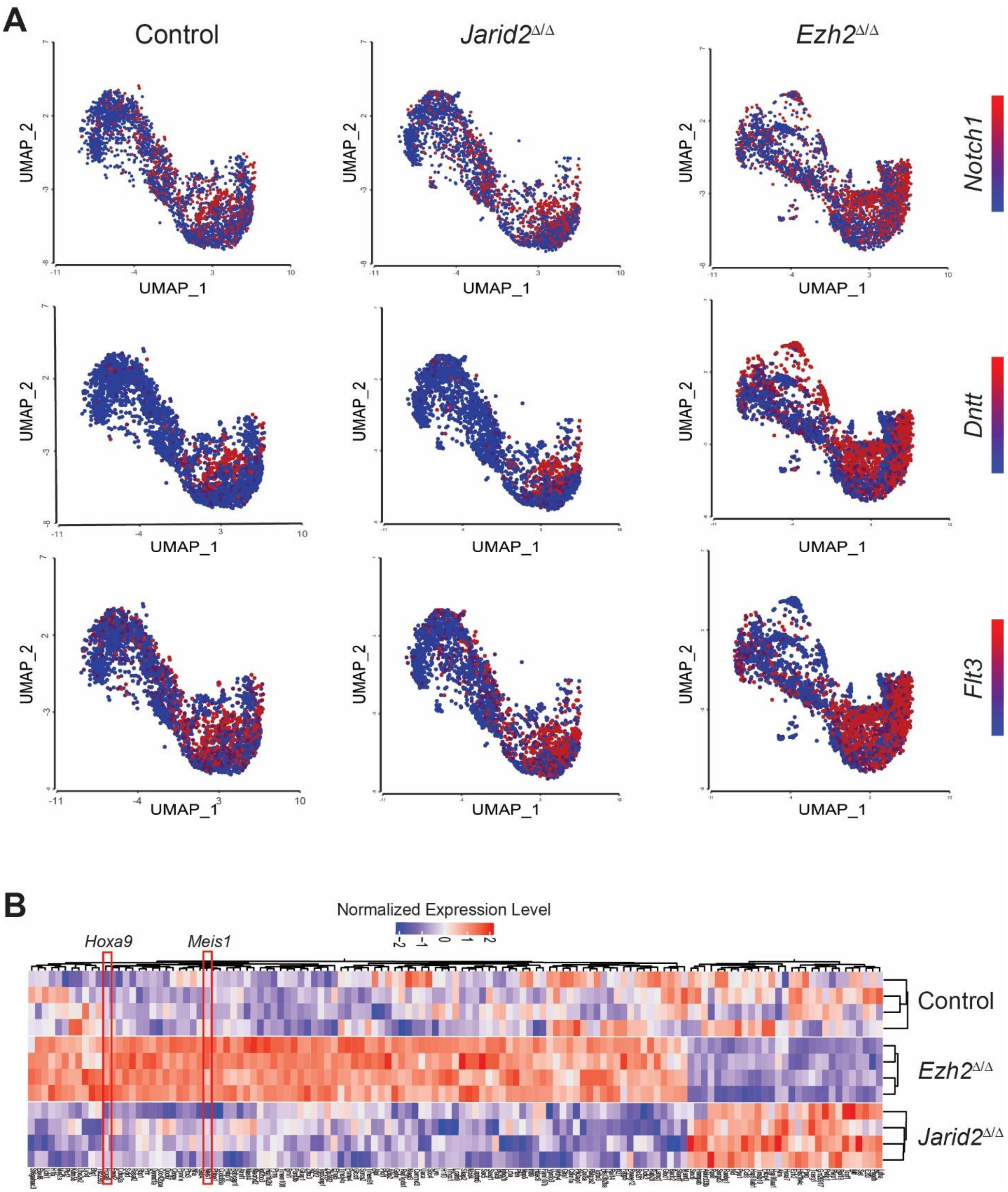


**Figure S1: Single cell gene expression analysis of HSPCs.**

A) UMAP expression plots of MPP4 marker genes *Notch1*, *Dntt* and *Flt3* in control, *Jarid2*$^{\Delta/\Delta}$ and *Ezh2*$^{\Delta/\Delta}$ HSPCs.

B) Pseudobulk differential gene expression analysis of MPP4 cluster cells showing upregulation of *Hoxa9* and *Meis1* in *Ezh2*$^{\Delta/\Delta}$ cells.

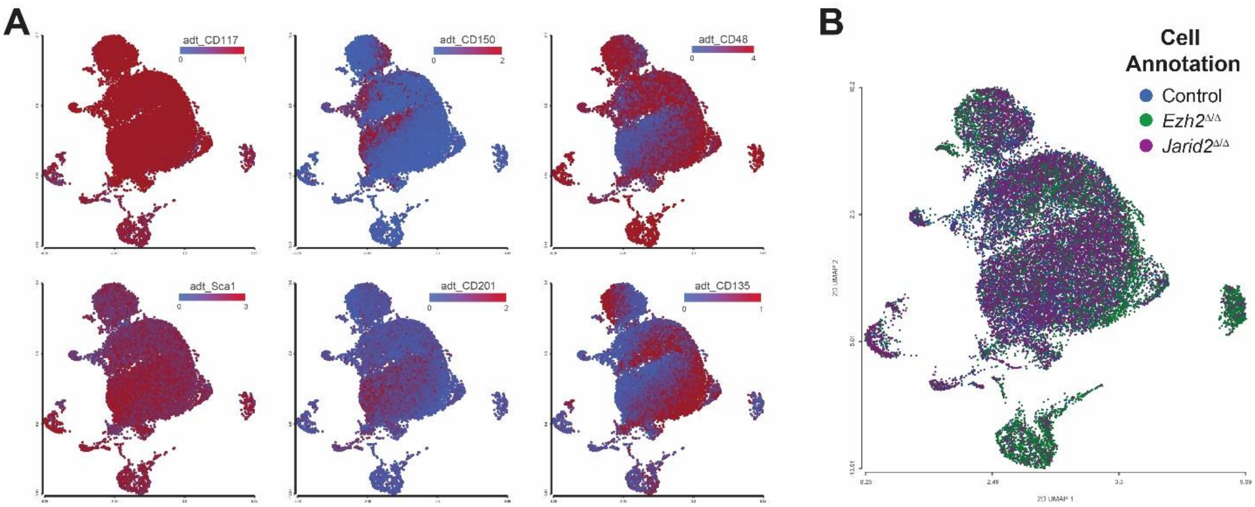


**Figure S2: Identification of HSPC clusters by antibody-derived tags in CITE-seq.**

A) Single cell UMAP showing expression of antibody-derived tags (adt) for cell surface markers CD117 (c-Kit), CD150, CD48, Sca-1, CD201, and CD135 (Flk2).

B) UMAP showing distribution of different genotype cells throughout the clusters with enrichment of *Ezh2*$^{\Delta/\Delta}$ cells in B-cell progenitor clusters.

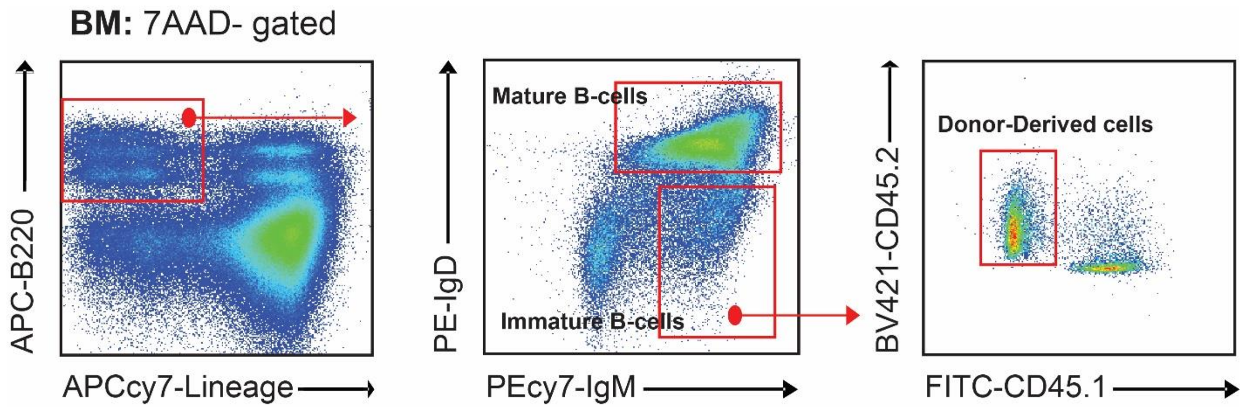


**Figure S3: Flow cytometric identification of B-cell populations.**

Representative flow cytometry gating strategy showing separation of donor-derived mature from immature B-cells in the bone marrow.
